# Insights into the metabolic specificities of pathogenic strains from the *Ralstonia solanacearum species complex*

**DOI:** 10.1101/2023.01.09.523232

**Authors:** Caroline Baroukh, Ludovic Cottret, Emma Pires, Rémi Peyraud, Alice Guidot, Stéphane Genin

## Abstract

All the strains grouped under the species *Ralstonia solanacearum* represent a species complex which collectively constitute a devastating plant pathogen responsible of many diseases on agricultural crops throughout the world. The strains have different lifestyles and host range. Here we sought whether specific metabolic pathways contribute to strain diversification. To this end, we carried out systematic comparisons, followed by manual expertise on 11 strains representing the diversity of the species complex. We reconstructed the metabolic network of each strain from its genome sequence and looked for the metabolic pathways differentiating the different reconstructed networks and, by extension, the different strains. Finally, we conducted an experimental validation by determining the metabolic profile of each strain with the Biolog technology, also in a comparative approach. Results revealed that the metabolism is conserved between strains, with a core-metabolism composed of 82% of the pan-reactome. The 3 species composing the species complex could be distinguished according to the presence/absence of some metabolic pathways, in particular one implying salicylic acid degradation. Phenotypic assays revealed that the trophic preferences on organic acids and several amino acids such as glutamine, glutamate, aspartate and asparagine are conserved between strains. Finally, the generation and assessment of the transcription factor *phcA* regulating virulence in each specie showed that the faster growth compared to the WT strain was conserved across *Ralstonia solanacearum* species complex.

**Author summary:** *Ralstonia solanacearum* is one of the most important threats to plant health worldwide, causing disease on a very large range of agricultural crops such as tomato or potato. Behind the *Ralstonia solanacearum* name are hundreds of strains with different host range and lifestyle, classified into three species. Studying the differences between strain allows to better apprehend the biology of the pathogen and the specificity of some strains. None of the published genomic comparative studies have focused on the metabolism of the strains so far. We developed a new bioinformatic pipeline to build high-quality metabolic networks and used a combination of metabolic modeling and high-throughput phenotypic Biolog microplates to look for the metabolic differences between 11 strains across the three species. Our study revealed that genes encoding for enzymes are overall conserved, with few variations between strains. However, at the level of the phenotype, more variations were observed. These variations probably result from regulation rather than the presence or absence of enzymes in the genome.

## Introduction

All the strains formerly grouped under the species *Ralstonia solanacearum* represent a species complex (abbreviated hereafter as RSSC) now comprising three distinct species, *R. solanacearum, R. pseudosolanacearum* and *R. syzygii* (1). These strains collectively constitute a devastating plant pathogen responsible of many diseases such as the bacterial wilt disease of solanaceous plants, the potato brown rot, the Moko disease on banana trees or Sumatra disease on clove (2, 3). Strains can infect over 250 hosts in 54 different botanical families (3), and are responsible of important economic loses throughout the world (4). The RSSC includes a large diversity of strains with phenotypic characteristics that may be specific for some strains (e.g. adaptation to cool temperatures, insect transmission…) and not necessarily related to phylogeny (e.g. host range). Historically, several classification systems have been used to differentiate these strains, either on host range (‘race’) or metabolic (‘biovar’) criteria (5) but these systems have been progressively abandoned as not robust enough (6). Based on genomic comparison methods, strains were classified in four phylogenetic groups called phylotypes (I to IV). Each phylotype corresponds roughly to the geographical origin of the strains but are not related to host specificity (7). The three distinct species of RSSC correspond to these phylogenetic groups : *Ralstonia pseudosolanacearum* corresponds to phylotype I and III, *Ralstonia solanacearum* corresponds to phylotype II and *Ralstonia syzygii* corresponds to phylotype IV, the latter being divided in three subspecies (1).

To better apprehend the genomic diversity of the *RSSC* strains, comparative genomic and proteomic analyses were performed on several strains either studying the whole genome (2, 3, 8–13) or focusing on specific genes such as Type 3 Secretion effectors (14) or antiphage systems (15). In particular, the species complex was shown to have a large pan-genome composed of at least 13128 distinct genes, and a core-genome composed of 3262 genes (13). The division into phylotypes of the RSSC is also supported by genomic and proteomic evidence (3, 13).

No study has attempted to connect possible metabolic specificities of strains with phenotypic life traits within the RSSC, an approach that has become possible with the advent of genomic-based methodologies. To date, the metabolism and trophic preferences (i.e. the identification of the metabolic substrates preferentially metabolized by a strain) were studied only in GMI1000 strain (16, 17). The metabolic network of *R. pseudosolanacearum* strain GMI1000 was reconstructed (16), which first provided a global view of the strains’ metabolism and catabolic capacities. This study revealed the existence of a metabolic trade-off between virulence functions and bacterial growth, with the cost of producing virulence factors reducing the maximum growth rate of the pathogen. It was shown that GMI1000 mutant strain disrupted from the central regulator PhcA are almost avirulent and could grow faster and on a wider range of substrates than the wild type strain (16). PhcA thus appears as a major regulator of both virulence and metabolism. Finally, the mapping of the substrates preferentially metabolized by the GMI1000 strain was carried out, thus allowing the identification of the compounds that sustain fast bacterial growth and were likely to be assimilated in tomato xylem sap (17).

In this manuscript, we present a systemic comparison of metabolism and trophic preferences of 11 strains belonging to all three species of the RSSC. To this end, the metabolic network of the 11 strains were reconstructed from their genome sequence, and trophic preferences of each strain was assessed using Biolog phenotypic microplates in order to establish the major convergences/divergences at this level. We also created *phcA* mutants in three strains to have a representative mutant of each species in order to establish the extent to which PhcA-dependent regulation of metabolism occurs within the species complex and whether the metabolic trade-off between virulence and growth observed in GMI1000 is conserved.

## Results

### Metabolic network reconstruction of 11 strains belonging to the three species

To represent the diversity of the RSSC, strains from each species of the species complex were chosen, with more strains *R. solanacearum* to better apprehend its diversity (Table 1). The genome of each strain was either taken from literature or sequenced by ourselves using PACBIO technology and structurally annotated (Table 1) using Prokka (18).

**Table 1.** List of the WT strains of the RSSC investigated in this study. The quality of each genome is shown in Fig S1. Number of proteins was computed from Prokka automatic annotation (18).

| Strain | Phylotype | Species | Plant origin | Country of origin | Genome sequence origin | Number of predicted proteins |
| --- | --- | --- | --- | --- | --- | --- |
| GMI1000 | I | *R. pseudosolanacearum* | Tomato | Guyana | (19) | 5060 |
| PSS4 | I | *R. pseudosolanacearum* | Tomato | Taiwan | This work | 5117 |
| RUN2340 | III | *R. pseudosolanacearum* | Potato | Madagascar | This work | 5119 |
| CFBP2957 | IIA | *R. solanacearum* | Tomato | French West Indies | This work | 4951 |
| K60 | IIA | *R. solanacearum* | Tomato | United States | (20) | 5081 |
| BA7 | IIA | *R. solanacearum* | Banana | Grenada | This work | 5035 |
| UW551 | IIB | *R. solanacearum* | Geranium | Kenya | (20) | 4755 |
| MOLK2 | IIB | *R. solanacearum* | Banana | Philippines | This work | 4868 |
| PSI07 | IV | *R. syzygii* | Tomato | Indonesia | (10) | 4812 |
| R24 | IV | *R. syzygii* | Clove | Indonesia | This paper | 4879 |
| BDBR229 | IV | *R. syzygii* | Banana | Indonesia | This paper | 4724 |

We developed an in-house reconstruction algorithm which we called Meroom (MEtabolic Reconstruction from Orthology and Ordered Metabolic models) to reconstruct automatically the metabolic network of each of the 11 strains. Briefly, this algorithm uses reference strains whose metabolic networks have either a high curation quality and/or are phylogenetically close to the organism whose metabolic network is to reconstruct. The first step of the algorithm consists in defining ortholog groups using Orthofinder (21). Then each metabolic reaction linked to an orthogroup is propagated into the metabolic network under reconstruction (Fig 1). The reference strains are ordered so that in case of conflicts the reference strain with the highest order of priority is trusted. This allowed to yield draft metabolic networks of high quality, with few false positives and few gaps. The closer the model strains are phylogenetically, the easier it is to reconstruct the metabolic networks to perform Flux Balance Analysis (22). Because the reference strain GMI1000 already had its metabolic network reconstructed (16), it allowed to generate in a straightforward manner the metabolic networks of the other 10 strains. Other bacteria such as *Cupriavidus necator* (23) or *E. coli* (24) were also used as models (see materials and methods for more details). The GMI1000 metabolic network was also reconstructed *de novo* with the Meroom algorithm to avoid any bias in the analysis. The GMI1000 network therefore refers to the new reconstructed metabolic network whereas the model network from Peyraud et al. (16) is referred to as ModelGMI1000. All the metabolic networks are available as File S1 in a table format and on a git repository in SBML formats (https://github.com/cbaroukh/rssc-metabolic-networks).

**Fig 1.**
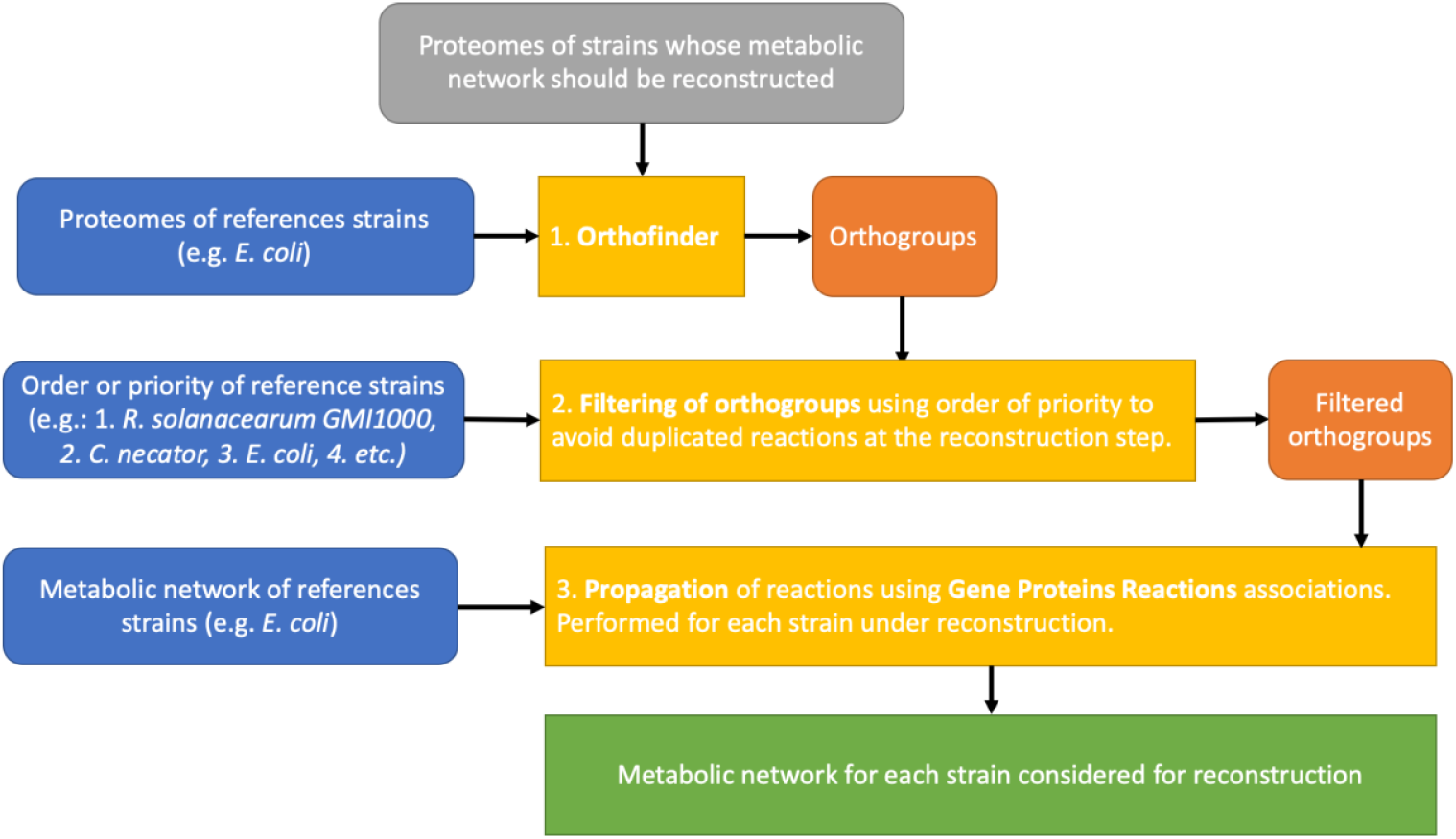
Meroom pipeline to reconstruct automatically draft metabolic networks of high quality. The algorithm relies on reference strains that are trusted for their metabolic network quality and/or which are phylogenetically close to the strains under reconstruction. The first step consists in using Orthofinder (21) to determine orthogroups between reference strains and strains under reconstruction. The second step consists in propagating reactions from reference metabolic models using gene-proteins-reactions associations and the orthogroups issued from step 1. The third step consists in merging the metabolic network obtained from each reference strain, for each strain under reconstruction. A more detailed description of the pipeline is available Fig S2.

The merged metabolism of all strains, i.e. the pan-metabolome, is composed of 2573 reactions and 2562 metabolites. The common metabolism of all strains, i.e. the core-metabolome, represents a big part of the pan-metabolome, since 2111 (82%) reactions and 2251 (88%) metabolites are present in all strains. A metabolic network contained in average 2390 reactions; the smallest one was obtained for BDBR229 (2315 reactions) and the largest for PSI07 and PSS4 (2440 reactions), both strains belonging to *R. syzygii* (Fig S3). Meroom propagates the complex links between genes and reactions formed by AND (protein complexes) and OR (isoenzymes). In the propagation, an orthologous gene participating in an enzymatic complex may be missing, so the link between genes and reactions is said to be incomplete. These reactions with incomplete gene links represent only 3.1 to 7.7 % of the reactions associated with a gene in the different networks.

### Tracking major metabolic differences among the strains

In order to identify metabolic markers that differentiate the 11 RSSC strains, we generated tables of presence/absence of the metabolic reactions for each strain and performed a clustering analysis and a Multiple Component Analysis (MCA) on these tables (Fig 2). The clustering analysis clustered strains belonging to the same phylotype (Fig S4), and strains belonging to same species. The MCA separated on the first axis (27% of explained variance) strains according to their phylotype, thus discriminating each species of the species complex. The second axis (21% of explained variance) separated strains within their phylotype. Phylotype and species could thus be recovered using only the presence/absence of reactions in each metabolic network.

**Fig 2.**
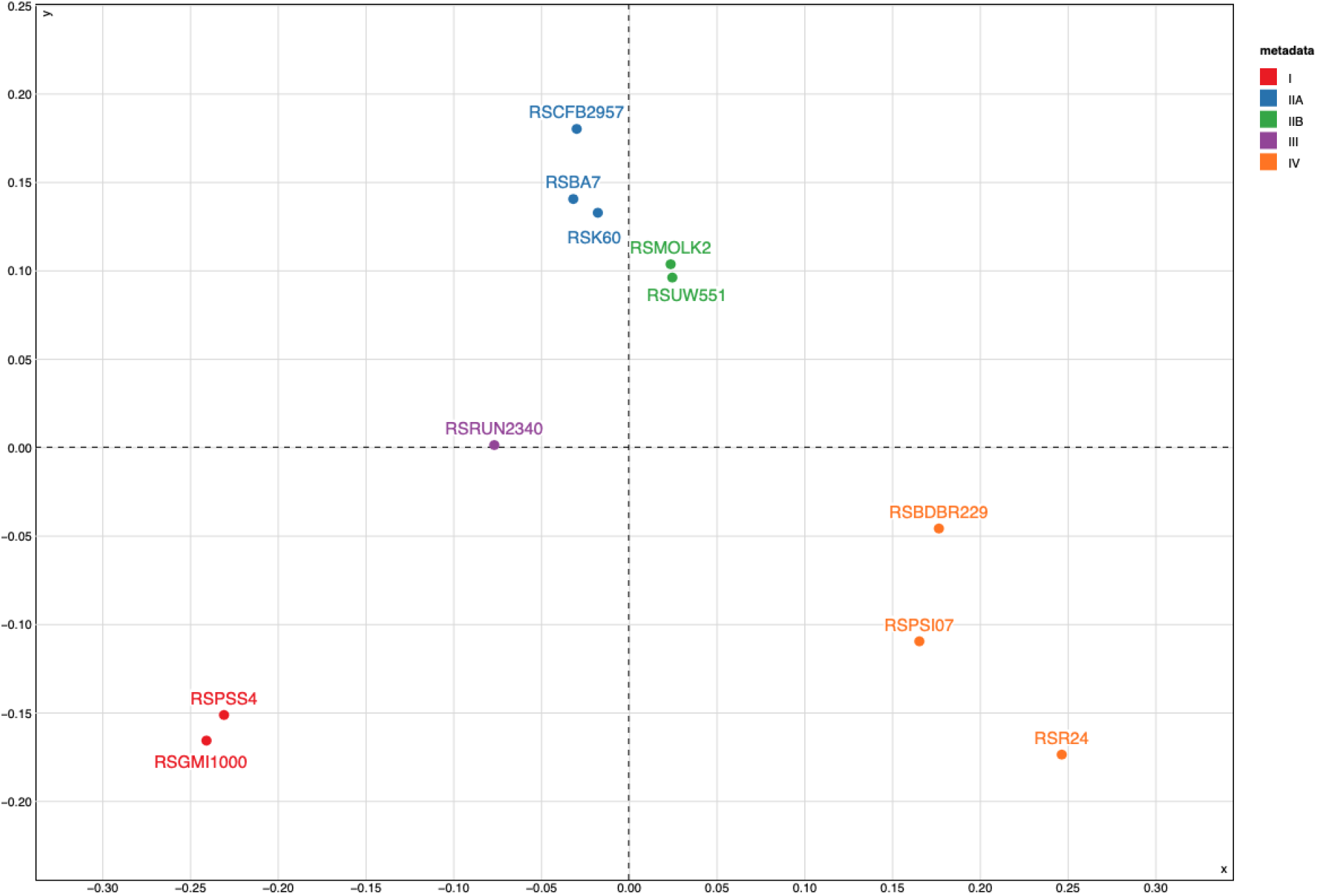
Multiple Component Analysis on the presence/absence of the reactions in the metabolic networks of 11 strains of *R. solanacearum*. Red: Phylotype I strains, Blue: Phylotype IIA strains, Green: Phylotype IIB strains, Purple: Phylotype III strains and Orange: Phylotype IV strains

From this MCA analysis, we sought to identify the metabolic pathways that distinguished the four phylotypes (see the list File S2). Looking into which reaction(s) contributed the most to the first axis, we found that several reactions belonged to the general pathway of benzoate degradation. Most of these reactions appeared in fact to be related to salicylate degradation pathways, showing that diverse pathways are able to degrade salicylate within the species complex (Fig. 3). Indeed, all strains possess a 4-aminobenzoate degradation pathway going through 3-oxoadipate and reaching the central core carbon network via succinyl-CoA and acetyl-CoA. All strains also have a salicylate degradation pathway via gentisate reaching the core metabolic network via fumarate and pyruvate. However, R24, K60 and CFBP2957 have lost some of the genes in the operon (2 to 4 genes) coding for this pathway (corresponding to RSc1085-RSc1091 in GMI100). Finally, all strains have a reaction converting salicylate to catechol (Fig 3). Phylotype I and phylotype III strains have another gentisate degradation pathway (RSc1821-1829 in GMI1000). Phylotype I has a catechol degradation pathway going through 2-oxopent-4-enoate to acetyl-CoA and pyruvate (Fig 3). All phylotype II and IV strains, except R24, have an operon degrading catechol to 3-oxodipate through cis,cis-muconate (Fig 3). In summary, according to the phylotype, different pathways are used by RSSC strains to degrade salicylate; only R24 appears to have a non-functional degradation pathway of this metabolite.

**Fig 3.**
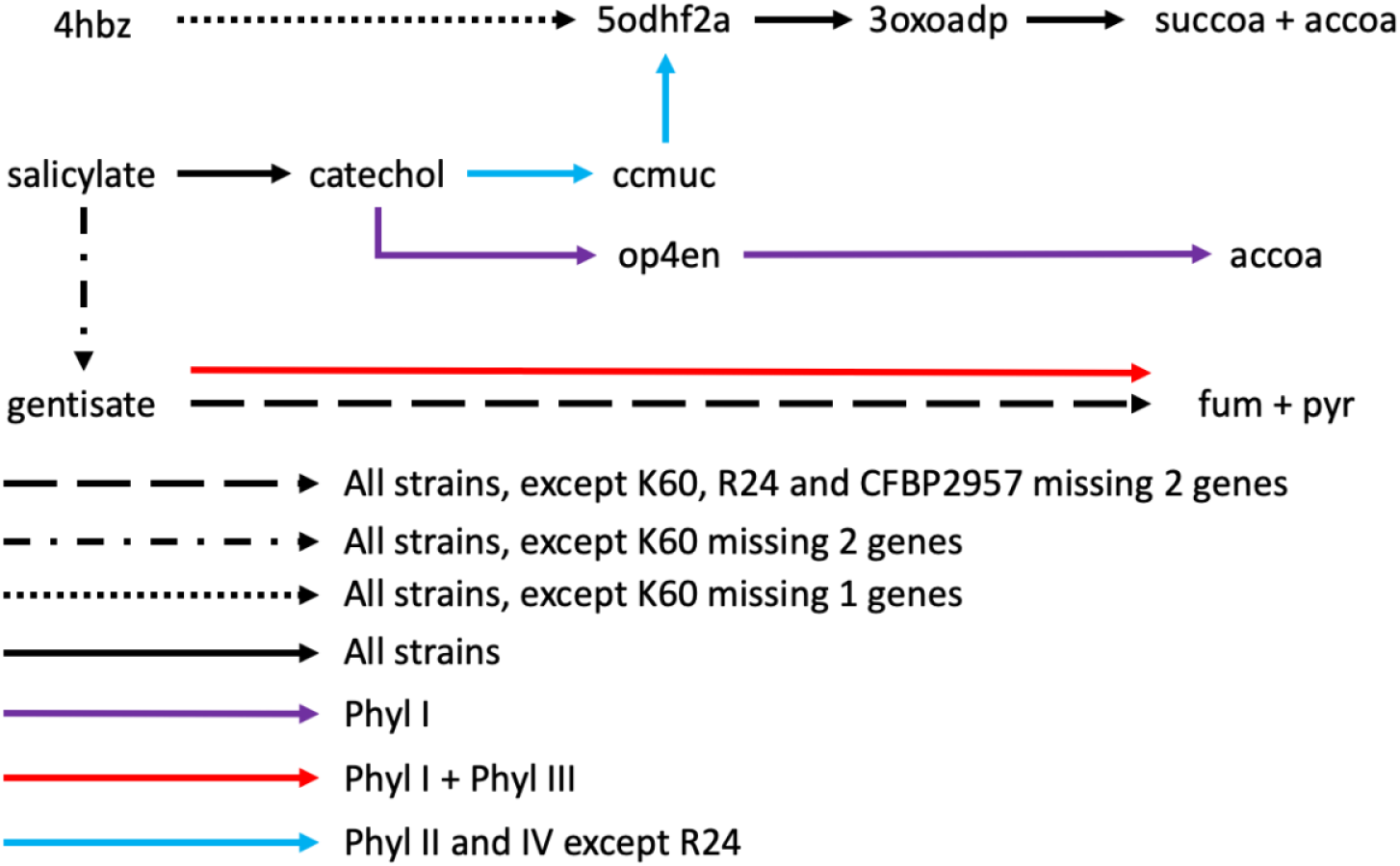
Different pathways to degrade 4-aminobenzoate and salicylate present in 11 strains. Black arrows: pathway present in the 11 strains investigated in this work. Dotted black arrow: present in all strains, K60 misses two genes catalyzing reactions in the pathway. Dashed black arrow: present in all strains, K60, R24 and CFBP2957 miss two genes catalyzing reactions in the pathway. Blue arrows: present in all phylotype II and IV strains except R24. Red arrow: present in all phylotype I and III strains. Purple arrow: present in all phylotype I strains. 4hbz: 4-aminobutyrate, 5odhf2a: 5-oxo-4,5-dihydrofuran-2-acetate, 3oxoadp: 3-oxoadipate, succoa: succinyl-coa, accoa: acetyl-coa, ccmuc: cis,cis-muconate, op4en: 2-oxopent-4-enoate, fum: fumarate, pyr: pyruvate.

Other differences contributing to the first axis of the MCA (listed File S2) relied on the presence/absence of several catabolic pathways, rather involving non-central metabolites, or single determinants in the primary metabolism which appear discriminating between groups of strains. For example, all the phylotype II and IV strains have a second glutamate dehydrogenase, converting glutamate to alpha-ketoglutarate, an extra phosphate import reaction, and a second pathway converting acetate into acetyl-CoA. Only phylotype IIB and IV strains possess a gallate degradation pathway. PSI07 (phylotype IV) and phylotype I and III strains have a degradation pathway for sarcosine, an intermediate metabolite for glycine-betaine degradation, and this pathway appears duplicated in phylotype I. Only phylotype I and III strains have a reaction converting N_2_O into N_2_ and a reaction repairing di-iron centers in Fe-S proteins, in agreement with previous reports (3, 25, 26), and phylotype IV strains have a thymine degradation pathway which is absent in other strains.

Looking at the second axis of the MCA, both R24 and BDBR229 miss an enzyme in ketogluconate catabolism, an enzyme involved in nitrate and nitrite import/export and 4 enzymes in the catabolism of glycogen. R24 also misses two additional enzymes belonging to the glycogen catabolic pathway, and one for glycogen synthesis (1,4 alpha-branching enzyme). Finally, three enzymes involved in galactonate degradation are missing in phylotype II strains. In conclusion, reactions discriminating strains are involved in salicylate degradation pathways, catabolism of specific substrates, reactions involved in secondary metabolism and some specific reactions involved in primary metabolism.

### Prediction of metabolic pathways critical for biomass growth in each reconstructed network

In order to predict *in silico* how the different strains were able to achieve growth under standard constraints, we performed Flux Balance Analysis (22) on each reconstructed network, by setting the same constraints previously used for strain GMI1000 (16). Briefly, growth was simulated on L-glutamate, with imposed excretion of putrescine, exopolysaccharides (EPS), ethylene, the diffusible signal molecule 3OH-PAME (methyl *3*-hydroxymyristate) and a protein substrate from the type II secretion system, as in Peyraud et al. (16). Unfortunately, for most strains, no growth and metabolite excretion were achievable, indicating that some essential reactions were missing in the corresponding reconstructed networks. To identify these essential reactions, assumed to be present in the ModelGMI1000, we have computed the systematic addition of each reactions from ModelGMI1000 missing in a given strain and then performed an *in-silico* reaction essentiality test on each of them. This allowed to unravel the missing reactions that were mandatory for growth and metabolites excretion in our FBA models. Detailed results are available in File S3.

First, BDBR229 missed nine reactions from the cobalamin synthesis pathway (vitamin B12), implying 8 enzymes belonging to the same operon. The entire operon has disappeared in BDBR229 (eq. RSp0614-RSp0628 in GMI1000). However, cobalamin is probably non-essential for growth (27) and contributes to a faster growth of the bacteria in media without the presence of cobalamin in the environment (28). In addition, BDBR229 has lost its S-adenosylmethionine decarboxylase (RSp1293 in GMI1000) necessary for synthesizing decarboxylated-S-adenosylmethionine, an intermediate metabolite which allows the synthesis of spermine and spermidine from putrescine. However, these polyamines are not always essentials for bacterial growth (29). Only putrescine, in GMI1000, was shown essential (30). Finally, R24 missed an 1,4 alpha glucan branching enzyme, necessary for the synthesis of glycogen. Looking in more details into the operon to which this enzyme belongs (RSp0235-RSp0242), the synteny of the operon is highly conserved in each strain, except for BDBR229 and R24. The operon is implied in glycogen synthesis and degradation. R24 misses 7 genes out of the 8 genes, and BDBR229 4 genes. The metabolic networks were modified to perform FBA for all strains by adding essential reactions or modifying the biomass equation. Results obtained also illustrates the high quality of the metabolic networks generated by Meroom since very few reactions (maximum 3) were modified or added in the metabolic network to be able to predict biomass growth and metabolite excretion. Some reactions were present in a strain but absent from ModelGMI1000. To know if the presence of these reactions conferred any gain in growth, we performed a similar analysis as for the missing reactions. The flux of each extra reaction was set to 0 to see if biomass growth was impacted. Results showed that none of the reactions conferred any significant gains in growth or metabolite excretion on glutamate (File S3).

### Prediction of metabolic robustness of RSSC strains through gene essentiality analysis

With the 11 metabolic networks reconstructed for each strain, we performed *in silico* a comparative gene essentiality analysis to estimate the metabolic robustness of each strain when growing on L-glutamate as sole carbon source (results listed in File S4). Beyond a group of genes (196) that are essential for all strains and belong to the essential central metabolism, this analysis identified 32 genes for which essentiality differed between strains (File S4). We examined this list in more detail to understand why these genes were predicted to be essential for growth in one strain and not in another, in order to uncover these specificities. The 32 genes were involved in majority in the synthesis of amino acids, or essential cofactors and vitamins such as folate, ubiquinone or flavin. Some of the genes were essential only in specific phylotypes as, for example, a reaction step in the synthesis of leucine which did not have any associated isoenzyme in phylotype II and IV strains, but had one in phylotype I and III strains (RSc1988 and RSp0329).

### Trophic preferences among the phylotypes

In parallel to the metabolic network analysis, the carbon and nitrogen trophic preferences of the 11 strains (Table 1) were studied using Biolog phenotypic microplates type PM1, PM2-A and PM3-B. Results revealed that BDBR229 has a fastidious growth compared to the other strains, since 192 hours instead of 96 hours were necessary to reveal a metabolic activity (Fig 4A). We developed a script to infer automatically if there was growth (resp. fast growth) and applied it on each substrate and for each strain (cf. material and methods for details, and File S5 for detailed results). Overall, there is a noticeable level of variation in metabolic versatility between strains, whether for nitrogen or for carbon sources. At both extremes, strain R24 appears to be able to metabolize twice as many substrates as either strain K60 or BDBR229 (101 versus 49 or 44, respectively). BDBR229 is an exception with a reduced versatility and only one substrate enabling rapid growth (Table 2), which probably explains the slow growth phenotype of this strain compared to the others. For K60, versatility is also reduced but seems to be compensated by a better ability to metabolize carbon substrates ensuring a rapid growth. All the strains could grow on 11 common carbon sources and 8 nitrogen sources (Table 2, File S5). When considering substrates which could sustain growth for 10 out of the 11 strains studied, we found 14 additional carbon sources and 6 additional nitrogen sources in common. The carbon sources included mainly organic acids and several amino acids but here too there were variations between strains, including strains within the same phylotype. This is particularly visible for certain amino acids (proline, alanine, serine, threonine) or sugars (sucrose, fructose, trehalose). We also detected automatically carbon substrates that could support a fast growth in at least 9 strains out of 10 (we excluded BDBR229, which does not have a ‘fast growth’ phenotype). We found that glutamine, glutamate, aspartate, asparagine, fumarate, citrate and malate could support a fast growth in most of the strains.

**Table 2.** Growth diversity of RSSC strains belonging to diverse phylotypes on amino acids, organic acids issued from the Krebs Cycle and sugars as carbon sources. The trophic preferences were assessed using Biolog phenotype microplates PM1 and PM2-A and an in-house script which detect automatically if there is growth (+) or fast growth (+++, see materials and methods for details). The rest of the substrates is available in File S5.

| Strain | GM1000 | PSS4 | RUN2340 | CFBP2957 | BA7 | K60 | UW551 | MOLK2 | Psi07 | R24 | BDBR229 |
| --- | --- | --- | --- | --- | --- | --- | --- | --- | --- | --- | --- |
| L-Aspartic acid | +++ | +++ | +++ | +++ | +++ | +++ | +++ | +++ | +++ | +++ | + |
| L-Glutamic acid | +++ | +++ | +++ | +++ | +++ | +++ | +++ | +++ | +++ | +++ | +++ |
| L-Asparagine | +++ | +++ | +++ | +++ | +++ | +++ | +++ | +++ | +++ | +++ | + |
| L-Glutamine | +++ | +++ | +++ | +++ | +++ | +++ | +++ | +++ | +++ | +++ | + |
| L-Alanine | + | + | +++ | + | +++ | - | + | + | + | +++ | + |
| L-Histidine | - | +++ | +++ | + | +++ | +++ | +++ | +++ | + | +++ | + |
| L-Serine | + | + | + | + | + | - | + | + | + | +++ | + |
| L-Threonine | - | + | +++ | + | +++ | - | +++ | + | - | - | - |
| L-Proline | - | - | +++ | - | +++ | +++ | +++ | + | - | +++ | + |
| L-Valine | - | - | - | - | + | + | + | + | - | + | - |
| L-Leucine | - | - | - | - | + | + | - | - | - | - | - |
| Succinic acid | + | + | +++ | + | +++ | +++ | +++ | + | + | +++ | + |
| Fumaric acid | +++ | +++ | +++ | +++ | +++ | +++ | +++ | +++ | +++ | +++ | + |
| L-Malic acid | +++ | +++ | +++ | +++ | +++ | +++ | +++ | +++ | +++ | +++ | + |
| α-Ketoglutaric acid | + | + | + | +++ | +++ | +++ | +++ | + | + | + | + |
| Citric acid | +++ | +++ | +++ | +++ | +++ | +++ | +++ | +++ | + | +++ | - |
| Pyruvic acid | +++ | +++ | +++ | +++ | +++ | - | +++ | +++ | +++ | +++ | + |
| α-D-Glucose | +++ | +++ | +++ | +++ | +++ | - | +++ | +++ | + | +++ | - |
| D-Trehalose | +++ | +++ | +++ | +++ | +++ | - | - | + | + | +++ | - |
| Sucrose | + | +++ | +++ | + | +++ | - | +++ | - | + | - | - |
| D-Fructose | - | + | + | - | - | - | + | + | + | +++ | - |
| D-Galactose | + | + | - | - | - | - | - | - | + | +++ | - |
| Phylotype | I | I | III | IIA | IIA | IIA | IIB | IIB | IV | IV | IV |

**Figure 4.**
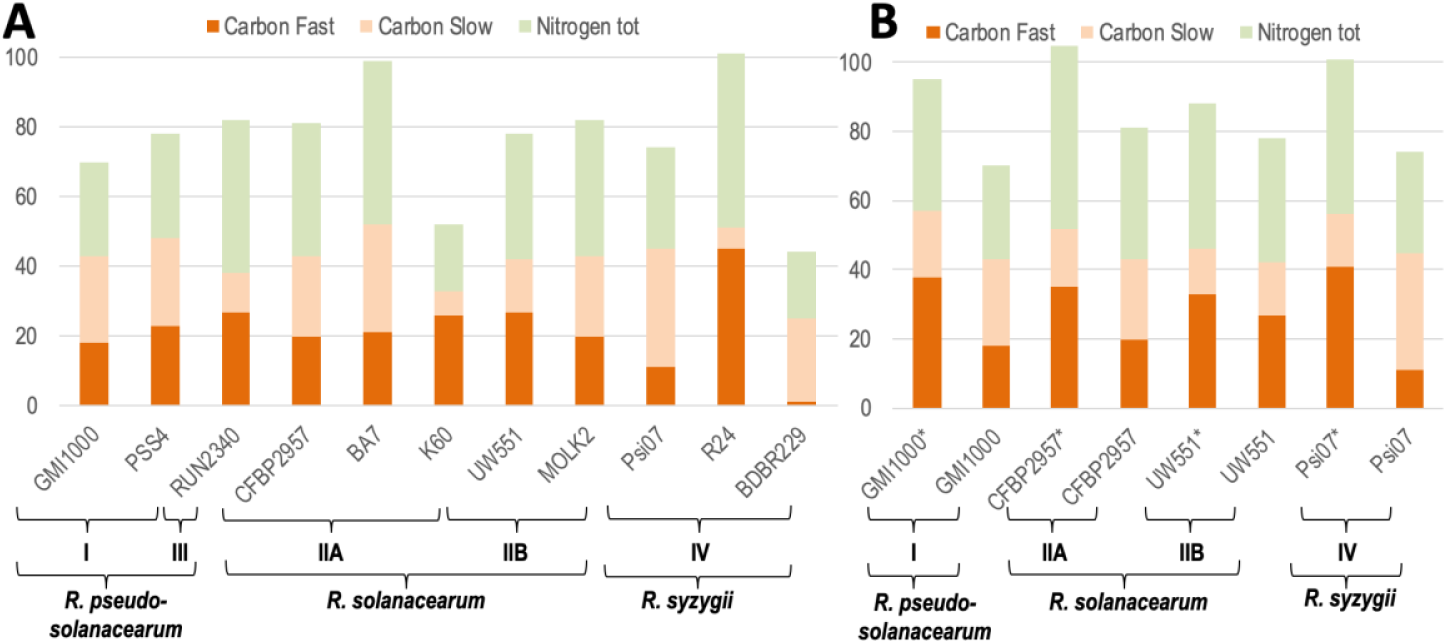
Number of carbon (resp. nitrogen) sources which can support growth for each wild-type strain (A) and *phcA* mutant strain (B). *: *phcA* mutant strain. Carbon sources were divided as sustaining a fast growth or a slow growth. Strains were grouped by phylotypes. The trophic preferences were assessed using Biolog phenotype microplates PM1, PM2-A and PM3-B and an in-house script which detect automatically if there is growth or fast growth (see materials and methods for details).

We performed a Principal Component Analysis (PCA) on the Biolog profiles of each strain using the maximal intensity (A) reached for each substrate as values characterizing each strain. The PCA could not distinguish the four phylotypes, contrary to the MCA performed on the metabolic network reconstructions (Fig S5). Similar results were obtained using the area under the curve (AUC) criterion instead of maximal intensity (data not shown). We performed hierarchical clustering, using the same table of values characterizing the strains (A and AUC). Here again, the strains were not clustered according to their phylogenetic relationships (Fig S6). Only strains from phylotype I clustered together when performing clustering on AUC for PM1 plate only (Fig S7). Specific markers of the relationship between these strains was the ability to metabolize sorbitol, dulcitol and mannitol. This finding confirmed previous results that identified a 22kb region specifically present in phylotype I strains (31) and was associated to the degradation of these three sugar alcohols.

### Conserved versus specific impact of the PhcA-dependent regulation on the metabolism of RSSC phylotypes

Since PhcA was shown to regulate metabolism (16), we sought to determine if this PhcA-mandated regulatory pattern on the pathogen’s metabolism was conserved within the RSSC. We therefore built *phcA* mutant strains in three strains representative of each species: CFBP2957 for *R. solanacearum* (phylotype IIA), UW551 for *R. solanacearum* (phylotype IIB) and Psi07 for *R. syzygii* (phylotype IV). We then performed Biolog phenotype microplates experiments for each of these mutants. In addition, we already had Biolog results for *phcA* mutant in GMI1000 (16). Results showed that any *phcA* mutant strain grew faster on more carbon substrates than the wild type, and could also grow on a larger number of substrates (Fig 4B). Overall, strains carrying the *phcA* mutation acquire a capacity to metabolize substrates ranging from 30 to 37% higher than the wild type except for strain UW551 where this rate is only 13%. This behavior reveals that, as in GMI1000 (16), PhcA exerts a catabolic repression in representative strains of the four phylotypes and so suggests the occurrence of similar growth/virulence trade-off. It is interesting to note that this catabolic repression takes place on a wide range of substrates, which again can vary between strains (Table 3). Several carbon sources that are not or poorly metabolized by wild-type strains can support a fast growth for *phcA* mutants in a majority of strains (proline, histidine, alanine, gluconate) while other carbon sources appear to be exploited more specifically by the *phcA* mutant of only a given strain (e.g. sucrose and malonic acid for strain CFBP2957). This observation underlines that a specific PhcA-mediated regulation may exist in some strains for some specific substrates.

**Table 3.** Comparison of fast growth between *phcA* mutants and WT strains on amino acids, organic acids issued from the Krebs Cycle, sugars and other discriminating metabolites as carbon sources. 2: both *phcA*-mutant and WT strains grow fast on the substrate. 1: Only *phcA*-mutant grows fast on the substrate. 0: neither the WT nor the *phcA-*mutant grow fast on the substrate. The trophic preferences were assessed using Biolog phenotype microplates PM1 and PM2-A and an in-house script which detect automatically if there is fast growth (see materials and methods for details). The rest of the substrates is available in File S5.

| Strain | Psi07 | CFBP2957 | UW551 | GMI1000 |
| --- | --- | --- | --- | --- |
| L-Aspartic acid | 2 | 2 | 2 | 2 |
| L-Glutamic acid | 2 | 2 | 2 | 2 |
| L-Glutamine | 2 | 2 | 2 | 2 |
| L-Asparagine | 2 | 2 | 2 | 2 |
| Fumaric acid | 2 | 2 | 2 | 2 |
| L-Malic acid | 2 | 2 | 2 | 2 |
| Pyruvic acid | 2 | 2 | 2 | 2 |
| α -D-Glucose | 1 | 2 | 2 | 2 |
| Citric acid | 1 | 2 | 2 | 2 |
| D-Saccharic acid | 1 | 2 | 2 | 2 |
| D-Trehalose | 1 | 2 | 0 | 2 |
| α -Ketoglutaric acid | 1 | 2 | 2 | 1 |
| Succinic acid | 1 | 1 | 2 | 1 |
| L-Proline | 1 | 1 | 2 | 1 |
| L-Histidine | 1 | 1 | 2 | 1 |
| Pectin | 1 | 2 | 1 | 1 |
| L-Alanine | 1 | 1 | 1 | 1 |
| Sucrose | 0 | 1 | 2 | 0 |
| L-Threonine | 0 | 1 | 2 | 1 |
| Acetic acid | 1 | 1 | 0 | 1 |
| D-Galactose | 1 | 1 | 0 | 1 |
| D-Fructose | 1 | 0 | 1 | 0 |
| L-Serine | 1 | 0 | 1 | 1 |
| D-Gluconic acid | 1 | 1 | 1 | 1 |
| m-Inositol | 1 | 2 | 0 | 1 |
| Butyric acid | 2 | 1 | 0 | 1 |
| Glycerol | 1 | 1 | 2 | 1 |

## Discussion

In this study, we set up a metabolic network propagation pipeline on RSSC strains using genomic sequences and reference networks including the manually curated network of GMI1000 strain. We sequenced (or re-sequenced) several strains in order to have at least three strains for each species of the RSSC with high quality genome sequence. This sample size is still limited and must weigh the generality of the conclusions, but the group of 11 strains used, beyond evolutionary (i.e. phylogenetic) relationship, also covers large phenotypic differences (broad versus narrow host range strains, adaptation to cool temperature, insect transmission versus root infection) and geographical origin.

Analysis of the 11 reconstructed networks reveals that the metabolic diversity is not so high between these strains, with a reactome (i.e. the set of possible metabolic reactions associated to the genome) comprising on average 2390 reactions, and a core-reactome consisting of 82% of the pan-reactome. This is in sharp contrast to the inter-strain comparison at the genomic level as the core-genome was estimated to comprise 1940 to 2370 genes (11.6% to 17.9% of the pan-genome, respectively), thus showing a high degree of genomic diversity between strains throughout the species complex (3, 12, 13, 32–34). The high proportion of the core-reactome thus reflects a very high (or even near complete) conservation of the core metabolism in the network of the different strains. Another point to consider is that most of the metabolic genes involved in secondary metabolism such as production of toxins or various uncharacterized diffusible molecules beside ralfuranones, Hrp-dependant diffusible factors, etc. (35–37) is also probably involved in adaptation processes and may vary between strains. However, the current state of knowledge of these secondary metabolic processes is relatively poor, thus making it nearly impossible to incorporate them into the reconstruction step of the metabolic networks.

Based on a criterion of presence/absence of reactions in the metabolic network of each studied strain, both the MCA and clustering analyses were congruent with phylogeny, distinguishing correctly in both cases the four phylotypes and three species. Strains from phylotype I and III clustered together, IIA / IIB strains were distinct from each other but clustered together in the phylotype II group, and phylotype IV strains also clustered even if they were the ones with the largest differences between strains (Fig2, Fig S4). This observation is in agreement with the current view of the phylogeny of the RSSC, with phylotypes I & III taxonomically closer together and grouped into the *R. pseudosolanacearum* novel species and phylotype IV in which a wider diversity is predicted (9, 13).

### No clear association between metabolic specificities and phenotypic traits but a common preference for organic acids and some amino acids

Apart from metabolic pathways that had already been identified as strain/phylotype-specific (metabolism of sugar alcohols (31) and nitrate assimilation (25)), the results point to a range of pathways involved in the catabolism of various ‘non-core’ compounds (salicylate, sarcosine, gallate, benzoate, galactonate…). Interestingly, gallate and salicylate are found abundantly in plants. In particular, salicylate, a plant molecule involved in defence against pathogens, was shown to be degraded by *R. solanacearum* to protect the bacteria against inhibitory levels upon infection (38). Our study reveals that the degradation of salicylate involves up to four distinct salicylic acid degradation pathways in the RSSC pan-reactome. Some of the salicylate degradation paths were only present in some phylotypes and not others, possibly reflecting a variety of evolutionary solutions to degrade this molecule, probably through selection for more efficient degradation in some strains. Intriguingly, only one strain (*R. syzygii* R24) appears to be unable to degrade salicylate, which raises questions about the existence of an as yet unidentified alternative pathway or a dependence on its particular lifestyle (insect-transmitted, and restricted to the clove tree host).

Two of the strains belonging to phylotype IV (BDBR229 and R24) have both very distinct behaviors from the other studied strains. Some metabolic determinants that are widely conserved in the species complex appear to be missing in these strains such as the glycogen degradation pathway, a functional salicylate degradation pathway for R24 or genes for the biosynthesis of the cofactor cobalamin in BDBR229. Biolog phenotyping confirmed that these two strains had atypical metabolic profiles, with BDBR229 having a limited number of substrates that could support growth, and R24, in contrast, being able to grow on the largest number of different substrates. These observations support the view that BDBR229 has a fastidious growth character, similar to other insect-borne plant pathogens (28, 39) but the case of R24, also insect-transmitted, remains enigmatic and it will probably be necessary to obtain other genomes of *R. syzygii* strains for a better understanding.

To a lesser extent, strain K60 (Phylotype II) also has characteristics that distinguish it from most other RSSC strains. K60 seems to be able to metabolize a smaller number of substrates than the average of the other strains but with greater efficiency (higher proportion of substrates promoting fast growth, see Table 2). The growth of this strain also appears to be deficient on several sugars (e.g. lacking some transporters such as for sucrose). It has recently been proposed that strain K60 may be representative of a new IIC clade (13) and it is therefore unclear at present whether the metabolic behavior of K60 is related to this phylogenetic distinction or not.

Beyond an apparent diversity in the trophic spectrum of the RSSC strains, our study highlights the importance of organic acids (such as citrate, malate or pyruvate) and amino acids (glutamine, glutamate, aspartate and asparagine) as a common substrate base supporting efficient bacterial growth in all strains tested (except BDBR229). Amino acids and sometimes organic acids are the main component of xylem sap of plants, with glutamine and asparagine often present in high amount in xylem sap (40). The experimental approach showed the importance of amino acids such as glutamine and asparagine in enabling abundant multiplication of strain GMI1000 in tomato xylem (17, 41). Amino acids and many organic acids are also present in soil extracts (42). The organic acids probably result from the decomposition of organic matter in the soil by bacteria and archaea, but also from root exudates (43). Roots also appear to exudate amino acids (43). Interestingly, L-malate was shown to be a chemoattractant of *R. solanacearum* (44) and it can be assumed that organic acids also play a crucial role in the survival of the bacteria in the soil.

### PhcA-mediated regulation reveals the metabolic potential of strains and also distinguishes substrate usage specificities

It is known that PhcA, a master transcriptional regulators in the RSSC, represses in a direct or indirect manner expression of some metabolic pathways since the corresponding mutants in strain GMI1000 could grow faster and on more substrates than the wild-type relative (16, 45). We generated *phcA* mutants in three additional strains, representing each species, including the IIA and IIB phylotype distinction. Characterization of the metabolic profile of these mutants showed that they could also grow faster and on more carbon and nitrogen substrates than wild-type strains, although there is significant variation in the number of additional carbon and nitrogen substrates between strains. Qualitatively, there were also significant variations in the type of substrates metabolized by the different phcA mutants, independent of phylogeny. These observations suggest that: (i) the trade-off between bacterial growth and virulence is globally conserved within the species, (ii) PhcA-mediated regulation can operate in a distinct manner on different metabolic genes depending on the strain, and that this regulation introduces a strong selectivity at the metabolic level, probably linked to the necessities of the strains’ life style and adaptation to their immediate environment.

The existence of such regulations could also explain why the PCA analysis of Biolog metabolic profiles does not overlap with phylogeny (unlike the analysis based on the presence/absence of reactions). Indeed, we can see that the presence of a metabolic gene does not necessarily imply its expression, and that the regulation that can take place will not depend on the phylogenetic origin of the strain. Another possible explanation to some discrepancies between Biolog profiling data and metabolic network prediction, is that the absence of a single gene such as a substrate transporter can lead to a negative phenotype even if the metabolic pathway is present (e.g UW551 misses the trehalose transporter).

## Materials and Methods

### Genome sequencing and annotation

Sequencing was performed as in Gopalan-Nair et al. (46) using Pacbio technology. Library preparation was performed at GeT-PlaGe core facility, INRAE Toulouse, France and SMRT sequencing at Gentyane core facility, INRAE Clermont-Ferrand, France. The mean reference genome coverage obtained was 227×.

For all the genomes, the Prokka software has been used to infer gene boundaries (18) with these parameters : --cdsrnaolap --coverage 60. The coding sequences were translated in amino acid sequences. The quality of each genome annotation has been measured with Busco (47), considering burkholderiale lineage as reference (Fig S1).

### Genome-scale metabolic network reconstruction

For the automatic reconstruction, ten metabolic models were used as reference (Table 4). The reference were chosen because they are using BiGG ontology (48), they are of high curation quality, they have a phylogenetic proximity to *R. solanacearum*, they were pathogens or had a similar lifestyle. Each model were downloaded in SBML format (49). Metabolic pathways information was added using BiGG database and ad-hoc scripts. Reaction and metabolite identifiers were standardized, based mainly on their formula, to avoid redundancies. For instance, two reactions involving exactly the same participants are identified by the same identifier.

**Table 4.** Reference metabolic networks used for the reconstruction step.

| Priority | Species | Lineage | Source of the metabolic model |
| --- | --- | --- | --- |
| 1 | *Ralstonia solanacearum* GMI1000 | Beta Proteobacteria | https://www.ebi.ac.uk/biomodels/MODEL1612020000 |
| 2 | *Cupriavidus necator* | Beta Proteobacteria | https://github.com/m-jahn/genome-scale-models/tree/master/Ralstonia_eutropha/sbml |
| 3 | *Escherichia coli* | Gamma Proteobacteria | http://bigg.ucsd.edu/models/iML1515 |
| 4 | *Xylella fastidiosa* | Gamma Proteobacteria | https://www.ebi.ac.uk/biomodels/MODEL2003100001 |
| 5 | *Xanthomonas oryzae* | Gamma Proteobacteria | https://www.ebi.ac.uk/biomodels/MODEL1912100001 |
| 6 | *Salmonella enterica* | Gamma Proteobacteria | http://bigg.ucsd.edu/models/STM_v1_0 |
| 7 | *Klebsiella pneumoniae* | Gamma Proteobacteria | http://bigg.ucsd.edu/models/iYL1228 |
| 8 | *Pseudomonas putida* | Gamma Proteobacteria | http://bigg.ucsd.edu/models/iJN1463 |
| 9 | *Bacillus subtilis* | Firmicutes | http://bigg.ucsd.edu/models/iYO844 |
| 10 | *Yersinia pestis* | Gamma<br>Proteobacteria | http://bigg.ucsd.edu/models/iPC815 |

We built a pipeline, called Meroom (MEtabolic Reconstruction from Orthology and Ordered Metabolic models) to build the metabolic network of each *R. solanacearum* strain studied. The Autograph method was used as starting point to infer the metabolic reactions (50). Autograph builds a new target metabolic network using only one reference network and performing orthology associations between the target and the reference genes. Gene-reaction associations are propagated from the reference to the target. Meroom improved the Autograph method by i) allowing several references at the same time and ii) allowing the reconstruction of several metabolic networks at the same time. In addition, Meroom orders the references so that the gene associations found in the first references are privileged over the others in case of conflict or duplication. It also computes the pan network and comparison matrices for reactions, metabolites, and pathways. Fig S2 gives an overview of the complete pipeline of Meroom approach.

Orthology relations between target and reference proteomes have been computed using Orthofinder (21). A percentage of 40% of identity has been set as parameter for the diamond execution. Then, orthologies are filtered according to the order of the references. For the first reference, all the orthology associations are kept. For the following references, orthology associations are kept only if the target gene has no ortholog in the previous references. For each target and for each reference, a target metabolic model is built. Then, the target networks obtained from each reference are merged into a unique target network. The reactions without gene-reaction association are kept only from the first reference. Finally, all the target networks built by Meroom are merged into a pan-network and comparison matrices are built (Fig S2).

Meroom uses met4j, a JAVA library for metabolic networks (http://metexplore.toulouse.inrae.fr/met4j), and is open-source (https://lipm-gitlab.toulouse.inra.fr/LIPM-BIOINFO/multiple-propagation). For convenience, a singularity package makes easier its installation and its usage (https://lipm-gitlab.toulouse.inra.fr/LIPM-BIOINFO/meroom-singularity).

### Computational simulations

Since some essential reactions for biomass production were missing in the metabolic network reconstructed by Meroom, we added manually these reactions to perform computational simulations. Tryptophan tRNA charging reaction was added because missing in the metabolic network of all strains. This was due to the fact that the GPR of this reaction relies only on an RNA sequence and not a protein, thus making it impossible to propagate to other networks, since orthologues were inferred at the protein level. Spermine was removed from the biomass equation of BDBR229 and CMR32. Cobalamin was removed from BDBR229 biomass equation. Glycolaldehyde dehydrogenase was added to all *R. syzygii* strains as well as K60 and RUN2340. 1,4-alpha-glucan branching enzyme was added in R24.

Simulations were performed using in-house Python 3.7 scripts. Reactions were parsed from metabolic network tabular files generated by Meroom. This allowed to create a numerical stoichiometric matrix and reversibility constraint vectors for each metabolic network. The linear programming solver CPLEX (Python API), developed by IBM, was used to solve the system and get solutions. All scripts and command lines are available online on GitHub: https://github.com/cbaroukh/rssc-metabolic-networks.

Flux balance analysis was performed using the following constraints: all lower bounds of transport reactions were set to 0 except water, hydrogen ion, potassium, phosphore, sodium, ammonium, sulfate, magnesium, chlore, iron, cobalt, manganese, molybdenum, oxygen and carbon dioxide. The others constraints used were the following : L-glutamate (R_EX_glu L_e set at -7.25 mmol.h^-1^.gDW^-1^), 3OHPAME (R_EX_3OHPAMES_e_ set at 1.5*10^−4^ mmol.h^-1^.gDW^-1^), EPS (R_EX_EPS_e_ set at 0.0062 mmol.h^-1^.gDW^-1^), putrescine (R_EX_ptrc_e set at 0.28 mmol.h^-1^.gDW^-1^), Tek (R_EX_Tek_e_ set at 2.7*10^−4^ mmol.h^-1^.gDW^-1^) and ethylene (R_EX_etle_e_ set at 0.129 mmol.h^-1^.gDW^-1^). Finally, non-growth associated maintenance (R_NGAME) was set at 8.38 mmol.h^-1^.gDW^-1^, and oxidation of Fe^2+^ to Fe^3+^ was set to 0 to avoid creation of energy from this reaction, which is not biologically relevant. Gene Deletion Studies were performed with similar constraints.

### Carbon substrate phenotyping

Phenotyping was performed using Biolog Phenotype Microarray plates PM1, PM2-A and PM3-B following the manufacturer’s protocol. An initial OD of 0.10 (600 nm) was used for inoculation. Incubation time was between 67 hours and 192 hours depending on the strain. Three independent replicates were performed.

Growth was assumed proportional to respiration and was assessed by calculating either the area under the curve (AUC) or the maximal intensity achieved (A, average on the top 10 values). Growth was considered effective if A > 50. Fast growth was considered if AUC > 7000. The raw Biolog results are available in File S6.

### Construction of *phcA* mutant strains

Disruptions of the *phcA* gene of the Psi07, CFBP2957 and UW551 receptor strains were created with the pGAΩ plasmid that was previously used to create the *phcA* mutant in strain GMI1000 (51). pGAΩ carries an insertion of the Ω interposon 255bp downstream of the *phcA* start codon. The Hin*dIII*-linearised construct was recombined in the genome of recipient strains through natural transformation (52). Competence of recipient strains has been achieved after growth for 48h in minimal medium (53) supplemented with 2% glycerol. Transformants were selected on medium supplemented with spectinomycin (40 µg/ml^-1^) and the genetic structure of the *phcA*::Ω recombinant locus was checked by PCR using the primers fw 5’-GGTACGACAACGAGTGG-3’ and rev 5’-TTCATCAGCGAGTTGACCGT-3’ (except for strain CFBP57: rev was 5’-TTCATCAGCGAATTGACCGT-3’).

## Supporting information

**File S1**. Metabolic networks of all 11 strains reconstructed as table format. The SBML format files are available in the github repository (https://github.com/cbaroukh/rssc-metabolic-networks).

**File S2**. Correlation scores of reactions contributing the most to the first two axes of the MCA.

**File S3:** Results of loss of functions and gain of functions of each reactions absent (resp. present) in the metabolic network of the strain but present (resp. absent) in the reference network of ModelGMI1000.

**File S4:** Gene deletion studies results for the 11 strains of *R. solanacearum* studied.

**File S5:** Growth and Fast growth of the strains on substrates for biolog microplates PM1, PM2-A and PM3-B

**File S6**. Experimental results of biolog microplates PM1, PM2-A and PM3-B for all 11 strains and 4 phcA mutants.

**Fig S1:** BUSCO Assessment results of the genome of the 11 strains of *R. solanacearum* studied.

**Fig S2:** Description of Meroom pipeline used to infer target metabolic networks

**Fig S3:** Size of the metabolic networks in terms of reactions.

**Fig S4:** Clustering tree on the presence/absence of metabolic reactions in the metabolic network of each *R. solanacearum* strain studied. Red: phylotype I. Purple: phylotype III. Blue: phylotype IIA. Green: phylotype IIB. Orange: phylotype IV.

**Fig S5:** Principal component analysis of results of Biolog profiles of the strains of *R. solanacearum* studied. Red: phylotype I. Pink: phylotype III. Blue: phylotype IIA. Green: phylotype IIB. Yellow: phylotype IV. PM1, PM2-A and PM3-B were used in triplicates to determine the profile of each strain. The maximal intensity (A) reached for each substrate was used as values characterizing each strain.

**Fig S6:** Hierarchical clustering based on results of Biolog profiles of the strains of *R. solanacearum* studied. Red: phylotype I. Pink: phylotype III. Blue: phylotype IIA. Green: phylotype IIB. Yellow: phylotype IV. PM1, PM2-A and PM3-B were used in triplicates to determine the profile of each strain. The maximal intensity (A) reached for each substrate was used as values characterizing each strain.

**Fig S7:** Hierarchical clustering based on results obtained on Biolog PM1-for the strains of *R. solanacearum* studied. Red: phylotype I. Pink: phylotype III. Blue: phylotype IIA. Green: phylotype IIB. Yellow: phylotype IV. PM1 were used in triplicates to determine the profile of each strain. The area under the curve (AUC) reached for each substrate was used as values characterizing each strain.

## Data Availability Statement

All data generated or analyzed during this study are included either in the manuscript in Figures, Tables or in Supplementary Data or in the github repository (https://github.com/cbaroukh/rssc-metabolic-networks).

## Model and Code Availability

Exploration, omics mapping and basic flux analyses can be performed on the model in MetExplore (54).

The main scripts and command lines used for the study are available on GitHub (https://github.com/cbaroukh/rssc-metabolic-networks).

## Author Contributions

Conceptualization: SG, CB, LC. Metabolic reconstruction and curation: LC, CB. Simulations and analysis: CB. Experiments: EP, RP, CB. Funding acquisition: AG, SG. Supervision: CB. Writing – original draft: CB, LC. Writing – review & editing: CB, SG, LC, RP, AG.

## Acknowledgements

The study was funded by the Centre National de la Recherche Scientifique, PEPs Modélisation de Processus Infectieux (2017-2019). Work in our laboratory is also supported by the French Laboratory of Excellence project “TULIP” (grant number ANR-10-LABX-41; ANR-11-IDEX-0002-02). The funders had no role in study design, data collection, and analysis, decision to publish, or preparation of the manuscript. We thank Toulouse Plant Microbe Phenotyping (TPMP) platform (Castanet-Tolosan, France) for the access to the Biolog phenotypic facilities. We also thank Patrick Barberis for his expert skills in the construction of the different mutant strains. We thank Ludovic Legrand for assembling the genomes of strains sequenced in the study.

