## Supplementary figures and images for "Insights into the metabolic specificities of pathogenic strains from the *Ralstonia solanacearum species complex*"

### Fig S

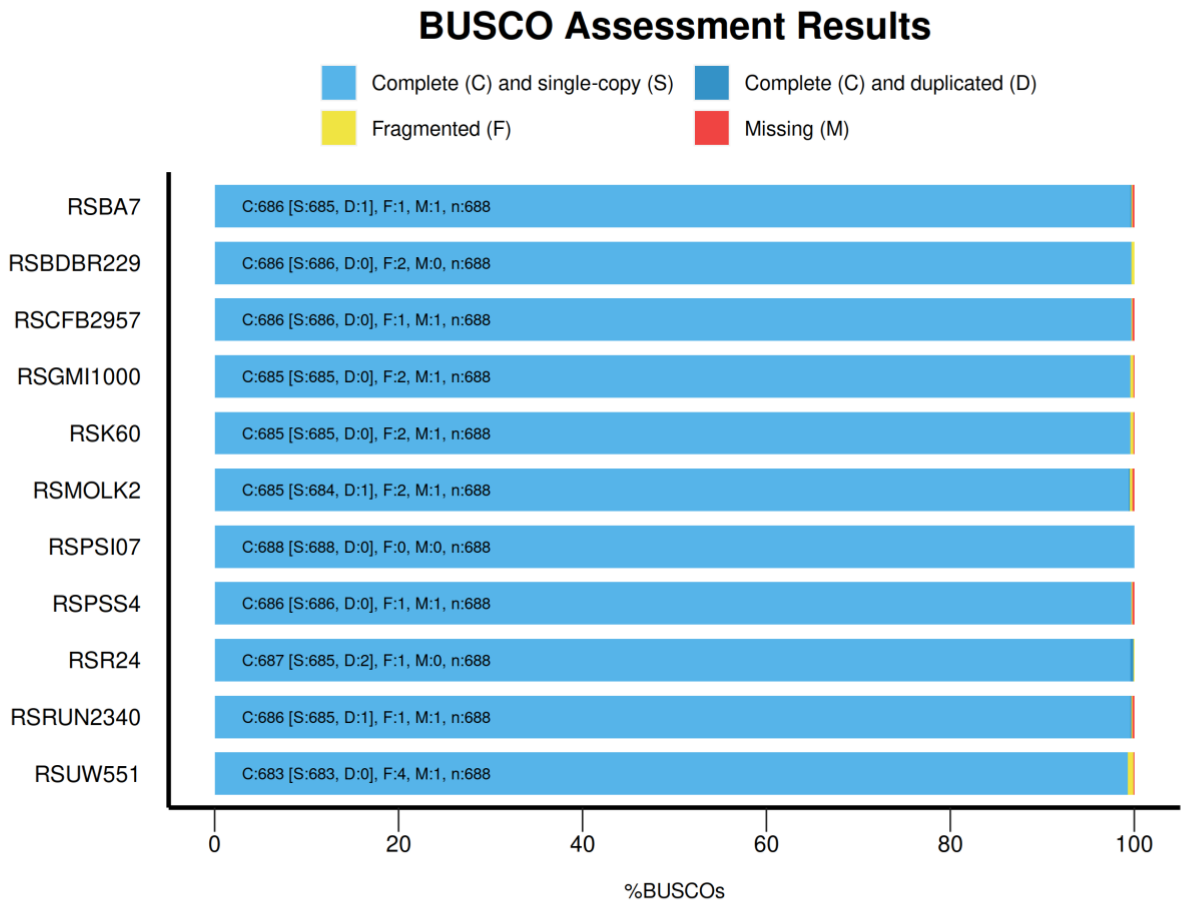

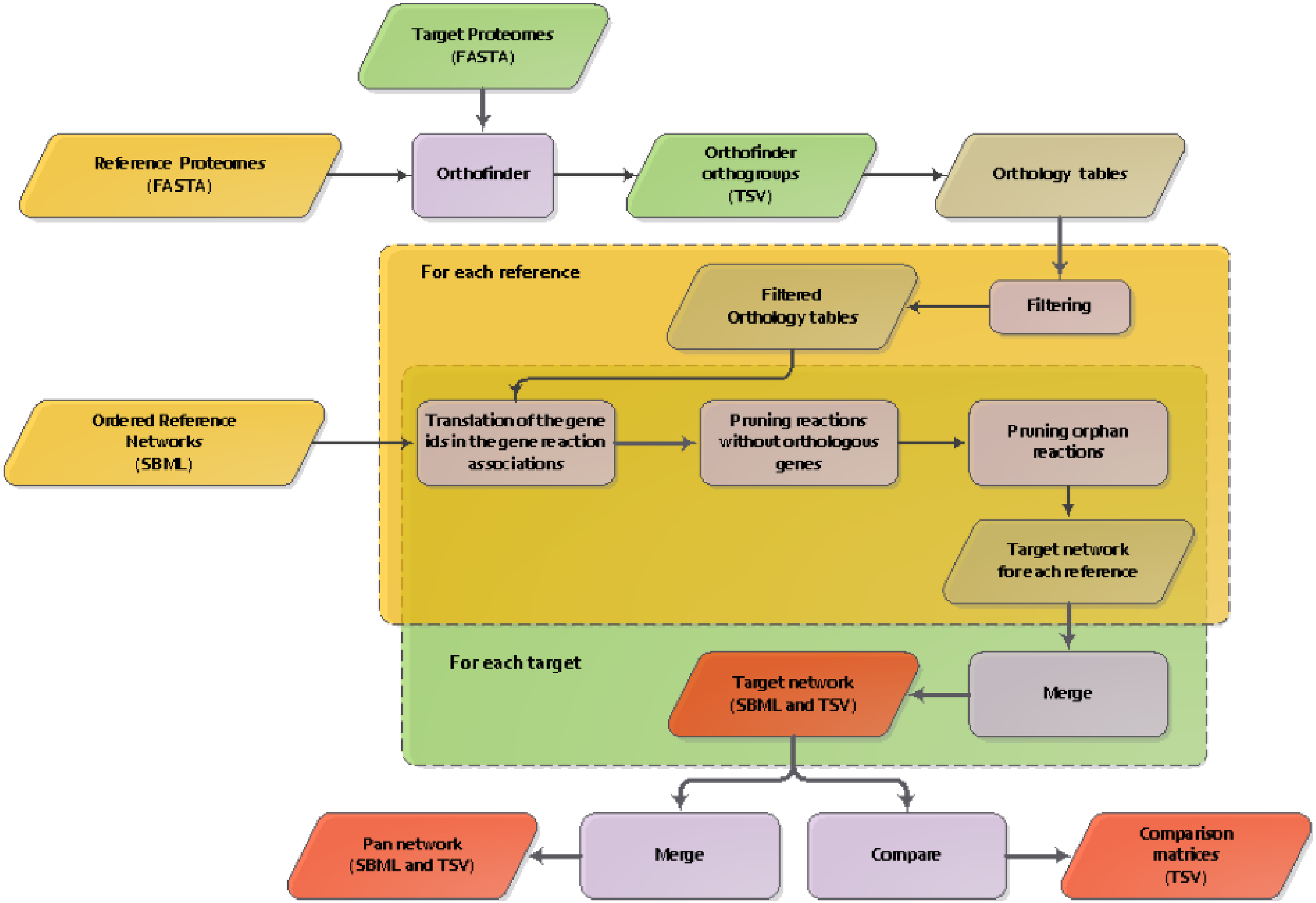

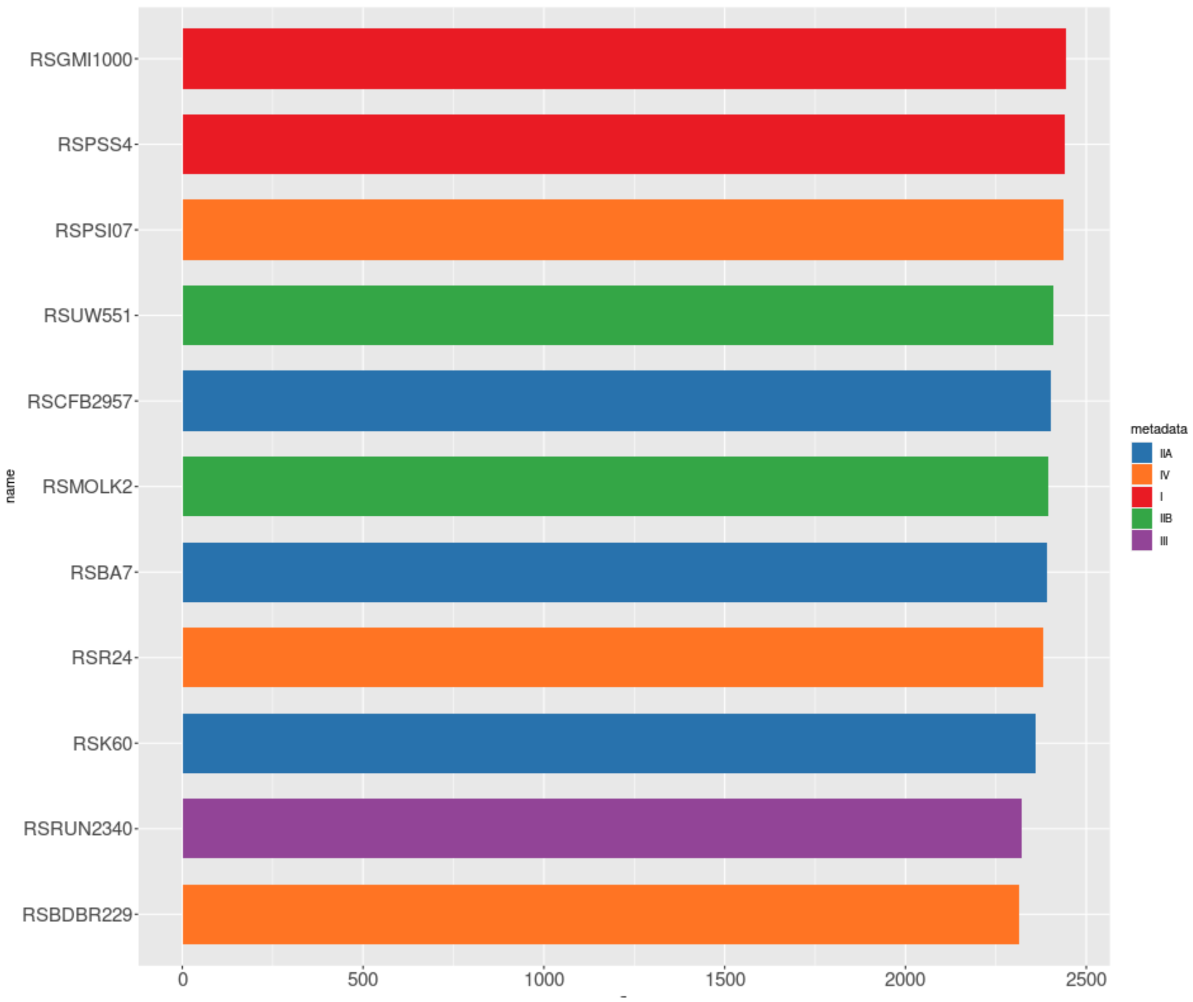

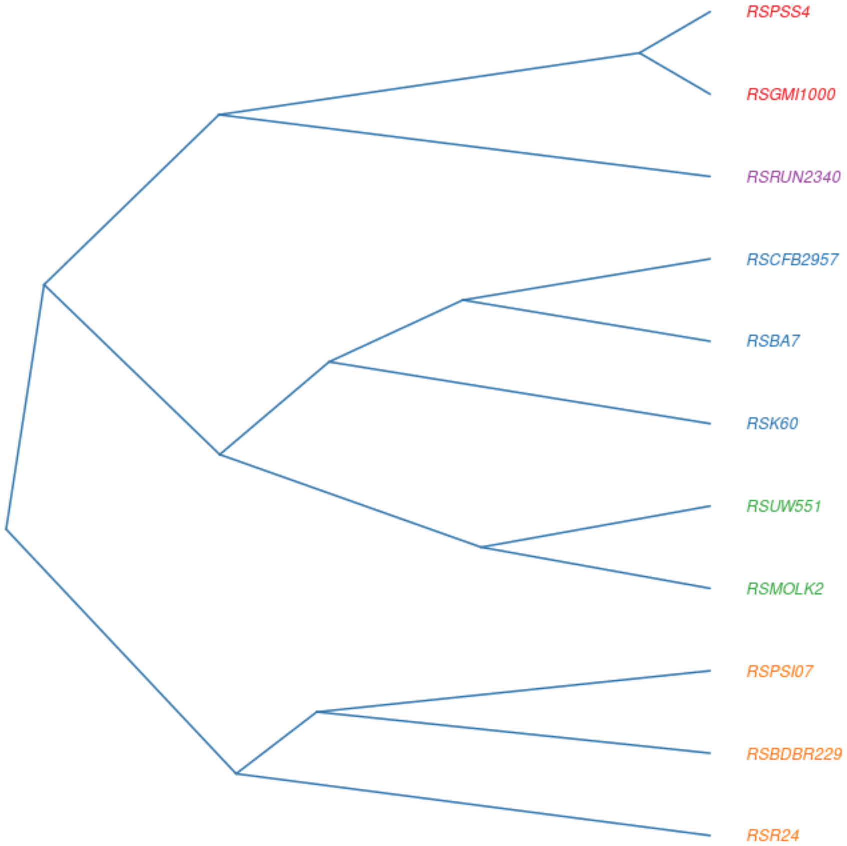

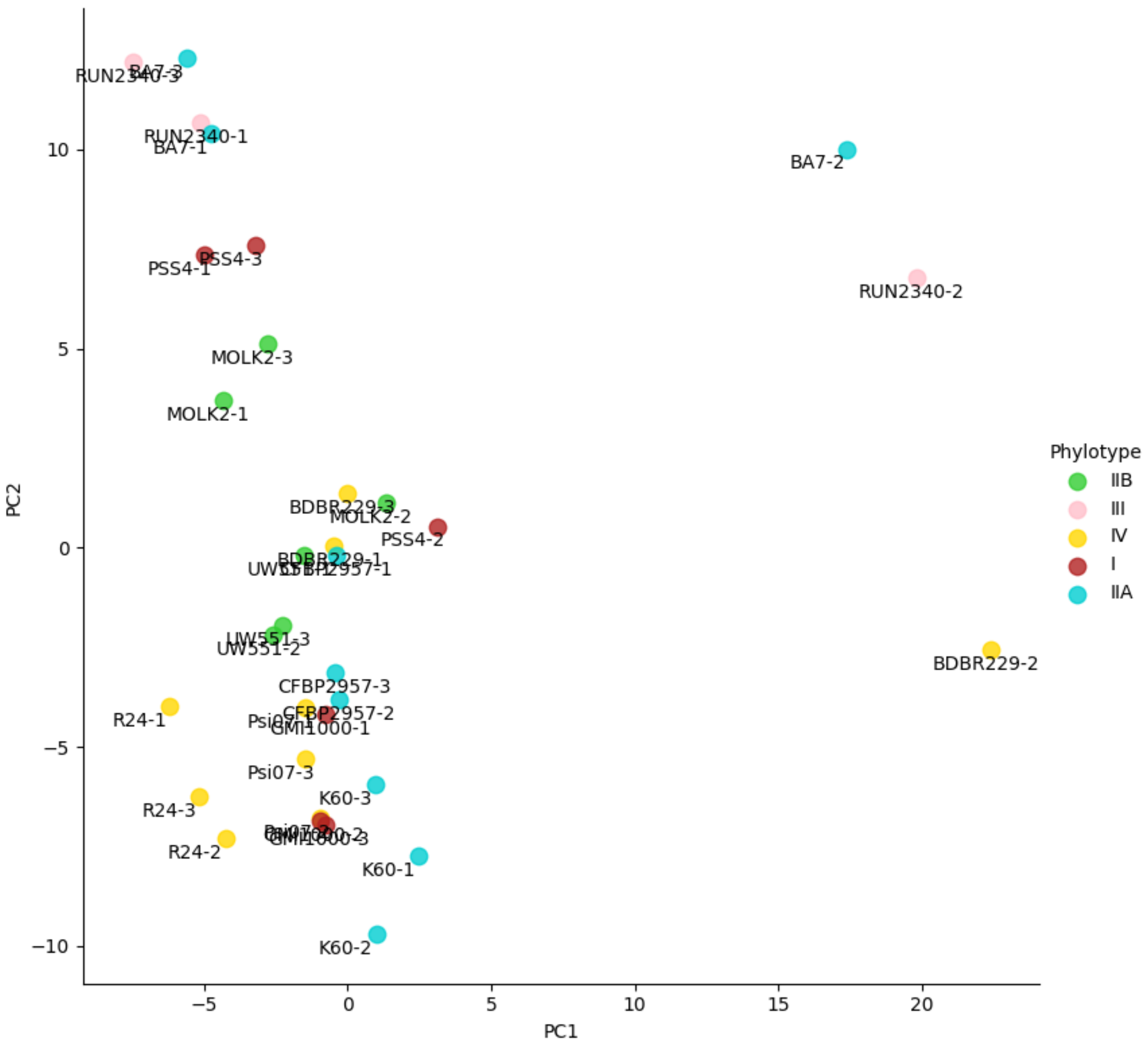

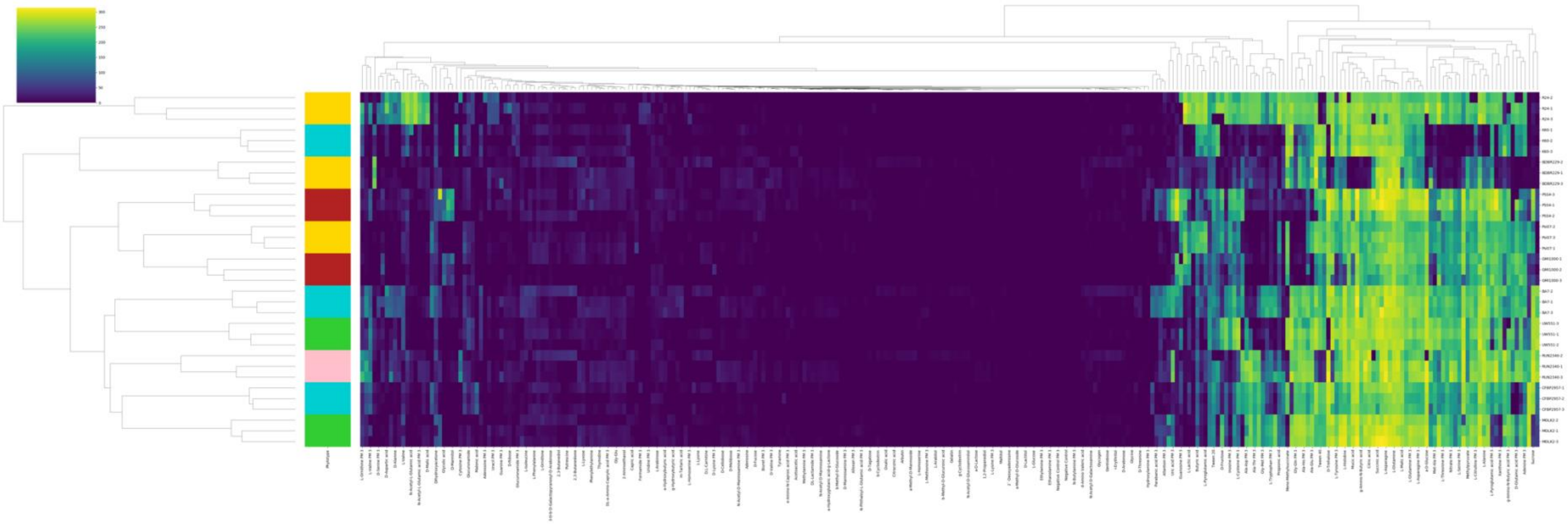

Fig S6

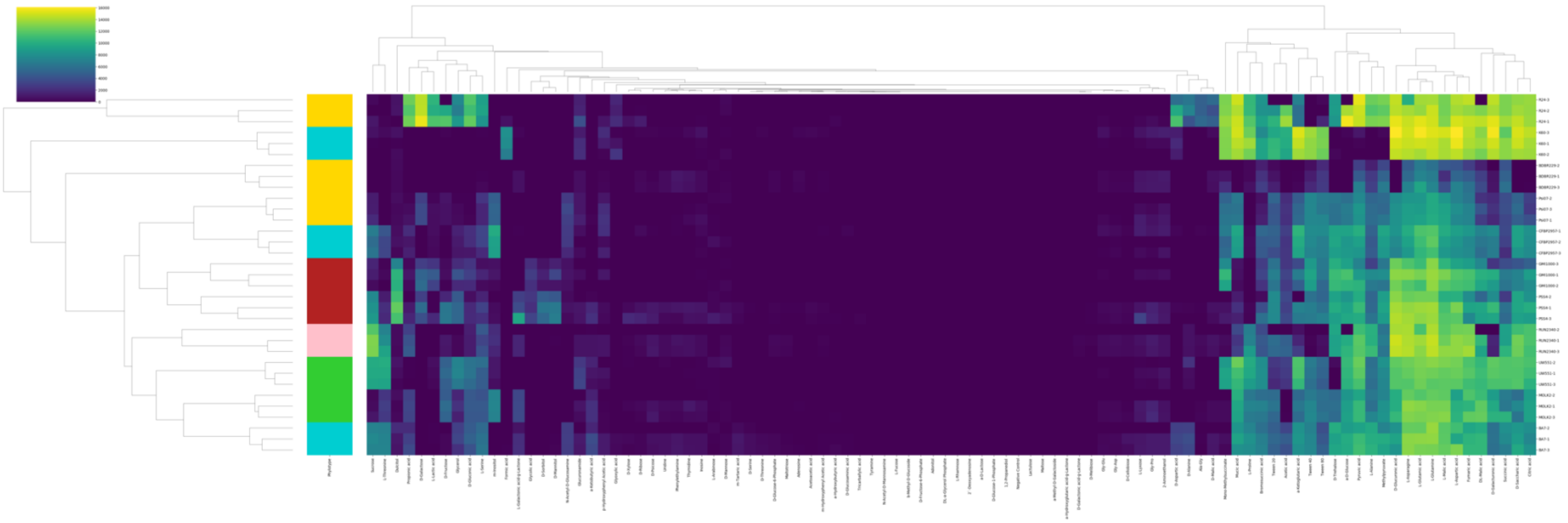

Fig S7
